# Antibody targeting of surface PSGL-1 glycoprotein leads to lymphoma apoptosis and tumorigenesis inhibition

**DOI:** 10.1101/2024.01.18.576249

**Authors:** João L. Pereira, Francisca Ferreira, Nuno R. dos Santos

**Affiliations:** i3S-Instituto de Investigação e Inovação em Saúde da Universidade do Porto; IPATIMUP-Institute of Molecular Pathology and Immunology of the University of Porto; FMUP-Faculty of Medicine of the University of Porto; ICBAS-School of Medicine and Biomedical Sciences, University of Porto; FEUP-Faculty of Engineering of the University of Porto, University of Porto

**Keywords:** Lymphoma, PSGL-1, Monoclonal antibody targeting, Apoptosis, Tumor growth

## Abstract

Lymphomas are a heterogeneous group of diseases that originate from T, B or natural killer (NK) cells. Lymphoma treatment is based on chemotherapy, radiotherapy, and monoclonal antibody (mAb) or other immunotherapies. The P-selectin glycoprotein ligand 1 (PSGL-1) is expressed at the surface of hematological malignant cells and has been shown to have a pro-oncogenic role in multiple myeloma and lymphoma. Here, we investigated the expression and therapeutic potential of PSGL-1 in T and B cell lymphomas. By flow cytometry analysis, we found that PSGL-1 was expressed in both T and B cell-derived lymphoma cell lines but generally at higher levels in T cell lymphoma cell lines. For most T and B cell-derived lymphoma cell lines, *in vitro* targeting with the PL1 mAb, which recognizes the PSGL-1 N-terminal extracellular region and blocks functional interactions with selectins, resulted in reduced cell viability. The PL1 mAb pro-apoptotic activity was shown to be dose-dependent, to be linked to increased ERK kinase phosphorylation, and to be dependent on the MAP kinase signaling pathway. Importantly, anti-PSGL-1 treatment of mice xenografted with the HUT-78 cutaneous T-cell lymphoma (CTCL) cell line resulted in decreased tumor growth, had no effect on *in vivo* proliferation, but increased the levels of apoptosis in tumors. Anti-PSGL-1 treatment of mice xenografted with a Burkitt lymphoma (BL) cell line that was resistant to anti-PSGL-1 treatment *in vitro*, had no impact on tumorigenesis. These findings show that PSGL-1 antibody targeting triggers lymphoma cell apoptosis and substantiates PSGL-1 as a potential target for lymphoma therapy.

## Introduction

Lymphomas are clonal malignancies commonly divided as Hodgkin lymphoma (HL), which is derived from B cells and characterized by an inflammatory microenvironment, or non-Hodgkin lymphoma (NHL), which is derived from T-, B- or NK cells.^1–5^ These malignancies are first-line treated with chemotherapy, radiotherapy, or mAbs targeting CD19, CD20 and other markers.^1,3,6,7^ About 60-70% of refractory/relapsed HL patients respond to immune checkpoint inhibitor treatments, possibly due to increased expression of PD-1 ligands and an inflammatory lymphoma microenvironment.^6,8–10^ However, NHL are often resistant to checkpoint inhibitor blockade therapies, with approximately 20-40% response obtained in clinical trials.^8,9,11^ Most NHL present a non-inflamed microenvironment with sparse immune infiltrates, and express transcriptional programs that promote immune evasion, such as the downregulation of antigen presentation.^9^ There is, therefore, a need for new strategies to improve lymphoma treatment.^12^

PSGL-1, encoded by the *SELPLG* gene, is a major ligand of P-, E- and L-selectin, not only widely expressed in normal lymphoid and myeloid cells but also expressed in hematological malignancies.^13–21^ PSGL-1 is normally involved in cell trafficking, rolling and tethering to blood vessels, as well as homing to secondary lymphoid organs and recruitment to inflammatory sites.^14,22–29^ Recently, PSGL-1 has been shown to bind to V-domain immunoglobulin suppressor of T cell activation (VISTA) under acidic conditions, typical of tumor microenvironments.^30^ It was reported that PSGL-1 was important for the generation and function of regulatory T cells and B cells.^31–33^ Furthermore, studies using *Selplg* knockout mice showed that PSGL-1 deficiency led to immune overactivation and triggered an autoimmune syndrome.^34^ In addition, PSGL-1-deficient mouse T cells showed prolonged T cell activation, enhanced T cell receptor (TCR) signaling, downregulation of inhibitory receptor expression and increased cytokine production when challenged with viral and tumor antigens.^35^

PSGL-1 was found to be highly expressed in multiple myeloma and responsible for regulating the homing of these cells to the bone marrow microenvironment.^36^ Moreover, PSGL-1 was shown to be involved in macrophage-mediated multiple myeloma resistance to chemotherapeutic drugs.^37^ In primary effusion lymphoma, the *SELPLG* gene was found to be amplified in 23% of cases and the PSGL-1 protein to be highly expressed in the majority of cases.^38^ In anaplastic large cell lymphoma (ALCL), PSGL-1 was found to be highly expressed and correlated with CD30 and TCR signaling genes and was put forward as a potential therapeutic target.^20^ Finally, in lymphoma mouse models PSGL-1 expression was found to promote lymphoma development and dissemination to distant organs.^39,40^ Taking into consideration these findings, we sought to assess whether PSGL-1 could be a therapeutic target in lymphoma. Here, we report PSGL-1 expression at the surface of several human lymphoma cell lines. Using the PL1 PSGL-1 mAb, we found that PSGL-1 targeting induced apoptosis of all T-cell derived cell lines tested and several B-cell derived cell lines *in vitro.* More importantly, the PL1 mAb impaired HUT-78 cell line-derived CTCL tumor growth in xenografted mice by increasing tumor cell apoptosis.

## Materials and Methods

### Cell lines and culture

The Raji and Daudi BL cell lines were provided by Alexandre M. Carmo (i3S, Porto, Portugal). The L428, KM-H2 and HDLM-2 HL cell lines were provided by Ana X. Carvalho (i3S, Porto). The HUT-78 CTCL cell line was provided by Neil D. Perkins (Newcastle University, UK). The OCI-LY3 (ACC 761), SU-DHL-4 (ACC 495), U2932 (ACC 633) and DOHH-2 (ACC 47) diffuse large B-cell lymphoma (DLBCL) cell lines; SU-DHL-1 (ACC 356) and L-82 (ACC 597) ALCL cell lines; and HH (ACC 707) CTCL cell line were purchased from the DSMZ repository (Braunschweig, Germany). All cell lines were cultured in suspension using RPMI 1640 medium (Gibco) supplemented with 10% heat-inactivated fetal bovine serum (FBS; Gibco), 100 U/ml penicillin-streptomycin (Gibco) and 1 mM L-Glutamine (Gibco), maintaining a density between 1 x 10^5^ and 1 x 10^6^ cells/ml, in a 5% CO_2_ humidified incubator at 37°C. Absence of *Mycoplasma* was confirmed by 16S rRNA amplification with MGSO and GPO-1 primers.^41^ For cell growth and viability assays, cells were seeded at 1 x 10^5^ and viable cells counted in a Neubauer hemocytometer 24 h later, using trypan blue as a viability exclusion marker. Cells were cultured with complete RPMI medium with or without anti-PSGL-1 PL1 hybridoma supernatant (1:10 dilution, approximately 5-10 µg/ml), anti-PAX5 PCRP-PAX5-1B monoclonal hybridoma supernatant (1:10 dilution, approximately 5-10 µg/ml) or purified KPL-1 anti-PSGL-1 mAb (cat. no. 328802, Biolegend). The PL1 hybridoma developed by McEver, R.P. and the PCRP-PAX5-1B7 anti-PAX5 deposited by the Common Fund – Protein Capture Reagents Program were obtained from the Developmental Studies Hybridoma Bank, created by the NICHD of the NIH and maintained at The University of Iowa, Department of Biology, Iowa City, IA 52242. MEK 1/2 inhibitor U0126 (cat. no. EI-282, Enzo Life Sciences) was used at 10 μM for the time-points described in the figures. For PL1 and U0126 treatment (5 min), all cells were put under starvation conditions (RPMI without supplementation) 2 h prior and during PL1 5 min stimulation.

### Primary human T cell isolation

Peripheral blood mononuclear cells (PBMCs) were obtained, with ethical approval by the CHUSJ Ethics Committee (Ref 398/2020), from buffy coats of adult healthy donors (aged 18 – 53 years) provided by *Banco de Sangue, Serviço de Imunohemoterapia, Centro Hospitalar Universitário São João* (CHUSJ), Porto. PBMCs were isolated by density-gradient separation using Ficoll Paque Plus (Cytiva). A red blood cell lysis buffer (90% 160 mM NH_4_Cl, 10% 170 mM Tris-HCl) was used to clear the pellet of remaining erythrocytes (5 min, 37°C). Cells were washed in phosphate-buffered saline (PBS). T cells were isolated using the MojoSort Human CD3 T Cell Isolation Kit (cat. no. 480021, Biolegend) and EasySep Magnet (cat. no. #18000, StemCell Technologies) according to the manufacturer’s instructions.

### Flow Cytometry

Cell lines were washed with FACS buffer (PBS with 3% FBS and 10 mM NaN_3_), centrifuged at 300 g, and resuspended in FACS buffer containing phycoerythrin (PE)-conjugated PSGL-1 antibody (clone KPL-1, cat. no. 328805, Biolegend). After incubation on ice for 45-60 min, cells were washed in FACS buffer. Samples were analyzed using BD Accuri C6 or BD FACSCanto II. For PL1 and KPL1 flow cytometry experiments, PL1 or KPL1 antibodies were incubated for 1 h followed by PE conjugated goat anti-mouse secondary antibody (cat. no. 405307, Biolegend) for another hour. For apoptosis detection, lymphoma cells were stained with 7-AAD Viability Staining Solution (cat. no. 420404, Biolegend) and Annexin V (cat. no. 640920, Biolegend) according to the manufacturer’s instructions. Cells were washed and fluorescent signals acquired using BD Accuri (lymphoma cell lines) or BD Canto II (healthy donor T and B cells). Data was analyzed using FlowJo software.

### Mouse experiments

*Rag2^-/-^Il2rg^-/-^* mice, provided by Nuno L. Alves (i3S, Porto), were bred and maintained at the i3S barrier animal facility under 12:12-hour light:dark cycles with food and water *ad libitum*. All experimental procedures were approved by the i3S ethics committee and Portuguese authorities (*Direção-Geral de Agricultura e Veterinária*) and followed recommendations from the European Commission (Directive 2010/63/UE) and Portuguese legislation (*Decreto-Lei* n°113/2013). For cell line xenografting, 10×10^6^ HUT-78 or Raji cells were injected subcutaneously (s.c.; interscapular region) into male *Rag2*^-/-^*Il2rg*^-/-^ mice. These were randomized into 2 groups and treated intraperitonally (i.p.) at days 1, 4, 7, 11 and 14 with 100 µg of either anti-PSGL-1 PL1 mAb, purified from hybridoma culture by Magellan Biologics & Consulting (Torres Novas, Portugal; purity of 97%, determined by size exclusion high-performance liquid chromatography; endotoxin level of 0.8 EU/mg, determined by LAL assay with Endosafe PTS, Charles River Laboratories) or mouse IgG (ChromPure Mouse IgG, cat. no. 015-000-003, Jackson ImmunoResearch). Tumor dimensions were measured with calipers at the indicated time-points. Mice were euthanized by CO_2_ inhalation 15 days post-injection (dpi; HUT-78-injected mice) or 36 dpi (Raji-injected mice).

### Immunohistochemistry

For immunohistochemistry, 4 µm tumor tissue sections were deparaffinized by immersion in xylene followed by ethanol. Antigen retrieval was performed with 1:100 dilution of Antigen Unmasking Solution, Citrate-based (cat. no. H-3300, Vector Laboratories) for 40 min at boiling temperature. Endogenous peroxidase was inactivated with 3% H_2_O_2_ in methanol at a 1:10 dilution. Nonspecific antibody binding was blocked using Ultravision Protein-block (Thermo Fisher Scientific). Next, tissue sections were incubated with rabbit anti-cleaved caspase 3 (1:200; cat. no. 9661, Cell Signaling Technology) or rabbit anti-Ki67 (1:500, cat. no. ab15580, Abcam). Dako REAL EnVision Detection System, Peroxidase/DAB+, Rabbit/Mouse (cat. no. K5007, Agilent) was used as secondary detection reagent. Images were obtained in a brightfield microscope Leica DM2000 LED using the software Leica Application Suite X (LAS X).

Percentage of cleaved caspase 3-positive lymphoma cells per tumor sample was evaluated in a blinded manner. Five images were taken representing five different areas of the tumor section and the percentage of cleaved caspase 3-positive cells was calculated using ImageJ software.

### Western blotting

Whole cell lysates from 2×10^6^ cells were prepared after two rounds of ice-cold PBS washing and resuspension in ice-cold RIPA buffer (10 mM Tris pH 7.4, 150 mM NaCl, 1 mM EDTA, 1% Triton X-100, 0.5% sodium deoxycholate, 0.1% SDS) with freshly added protease inhibitors (10 μg/ml aprotinin, 10 μg/ml leupeptin and 1 mM phenylmethylsulfonyl fluoride) and phosphatase inhibitors (10 mM 4-nitrophenyl phosphate disodium salt hexahydrate, 15 mM β-glycerophosphate disodium salt hydrate and 1 mM Na_3_VO_4_) for 10 min. After 16,000 g centrifugation, the total protein concentration of each sample’s supernatant was quantified with DC Protein Assay (cat no. 5000116, BioRad) in a 96-well plate. Then, samples were denatured using sodium dodecyl sulphate (SDS) sample buffer (62.5 mM Tris pH 6.8, 20% glycerol, 2% SDS and 5% β-mercaptoethanol) and heated for 5 min at 96°C. Protein samples were subjected to 8-12% SDS-polyacrylamide gel electrophoresis, together with PageRuler Plus prestained protein ladder (cat. no. 26619, Thermo Fisher Scientific) and transferred to a nitrocellulose membrane (cat. no. 10600009, GE Healthcare). Blotting efficiency was assessed by Ponceau membrane staining (P7170, Sigma-Aldrich). Nonspecific proteins were blocked with 5% nonfat dried milk in PBS with 0.1% Tween 20 and then stained with ERK2 (1:5000 dilution; cat. no. sc-154, Santa Cruz Biotechnology) and α-Tubulin (1:1000, cat. no. T6199, Sigma-Aldrich). Diluted 1:5000 horseradish peroxidase (HRP)-conjugated goat anti-mouse IgG (cat no. A00160; GenScript) and diluted 1:5000 HRP-conjugated goat anti-rabbit IgG (cat no. A00098, GenScript) were used as secondary antibodies. For phosphorylated-ERK (1:1000 dilution; cat. no. 9101S, Cell Signaling) hybridization, Tris-buffered saline with 0.1% Tween 20 was used for washing and antibody incubation as well as membrane blocking with 5% bovine serum albumin (cat. no. MB04602, NZYTech). The chemiluminescent signals were acquired by incubating the membrane with SuperSignal West Femto Maximum Sensitivity Substrate (cat. no. 34094, Thermo Fisher Scientific) and captured by ChemiDoc MP Imaging System (Bio-Rad). For membrane re-hybridization, an antibody stripping solution (10% methanol 10% acetic acid in distilled water) was used.

## Results

### PSGL-1 is expressed at the surface of human lymphoma cell lines

We aimed to determine the surface PSGL-1 expression levels in cell lines of different lymphoma subtypes. Flow cytometry analyses using a fluorochrome-conjugated KPL-1 mAb revealed that surface PSGL-1 was expressed in all cell lines, but with higher expression in T lineage CTCL (HUT-78 and HH) and ALCL (L-82, SU-DHL-1) cell lines, and lower expression in B lineage cell lines, especially in DLBCL cell lines (OCI-LY3, SU-DHL-4 and DOHH-2; Fig. 1A). The higher levels of PSGL-1 surface expression in T lineage as compared to most B lineage lymphomas might reflect the reported higher PSGL-1 expression in normal T cells as compared to normal B cells (Fig. 1B).^42^

**Figure 1.**
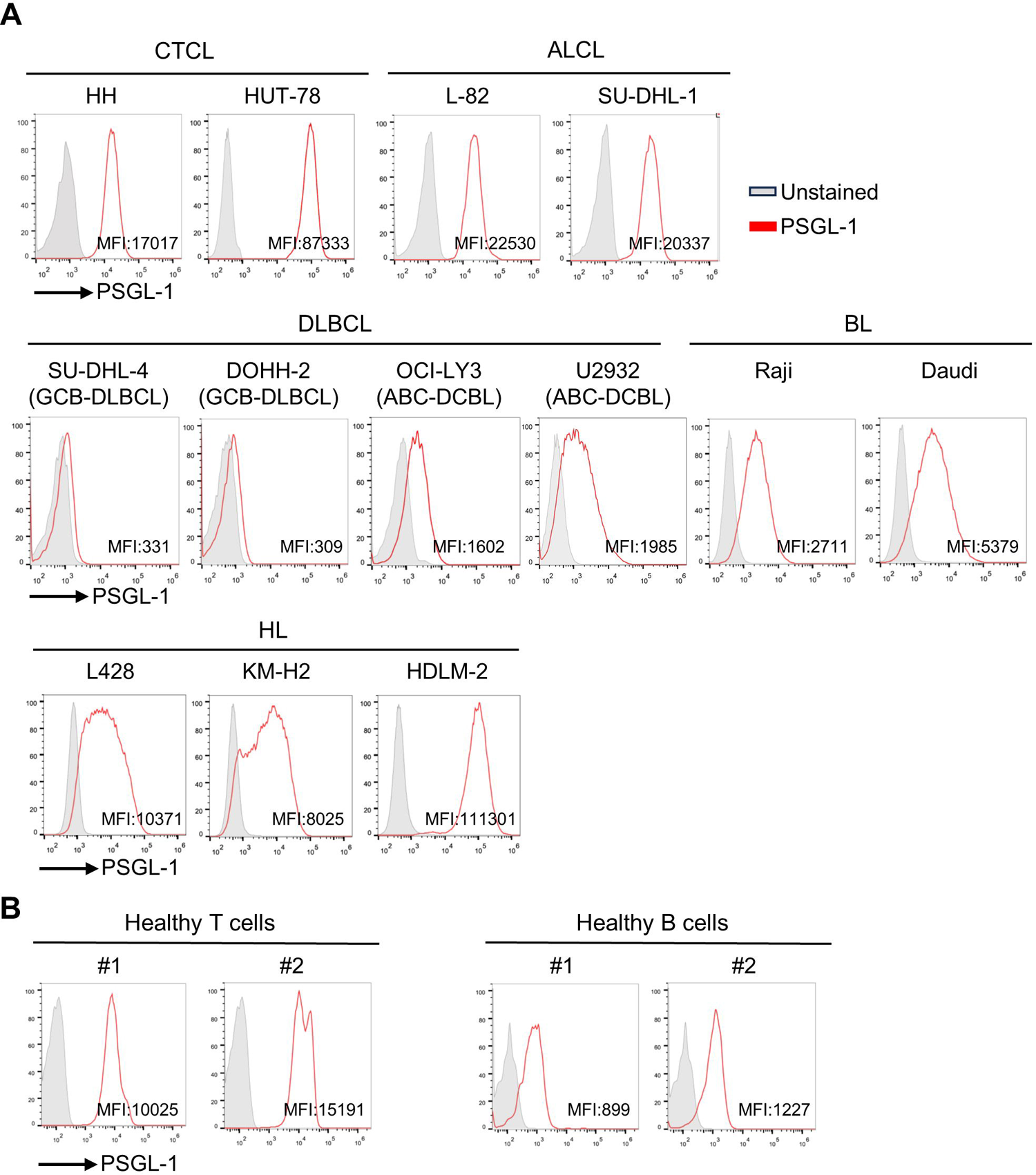
PSGL-1 is expressed in several human lymphoma cell lines. A) Flow cytometry histograms of surface PSGL-1 expression in human lymphoma cell lines detected in a BD Accuri analyzer. CTCL, Cutaneous T-cell lymphoma; ALCL, Anaplastic large cell lymphoma; DLBCL, diffuse large B-cell lymphoma; GCB-DLBCL, germinal center B DLBCL; ABC-DCBL, activated B-cell DLBCL; BL, Burkitt lymphoma; HL, Hodgkin lymphoma. B) Flow cytometry histograms of surface PSGL-1 expression detected in BD Canto II analyzer in two healthy donor (#1 and #2) peripheral blood T and B cells, after gating for CD3^+^ and CD19^+^, respectively.

### Lymphoma cell line *in vitro* treatment with a PSGL-1 antibody reduces viability and induces apoptosis

Given that PSGL-1 is expressed at the surface of many lymphoma cell lines, we hypothesized that antibody targeting of PSGL-1 could impair lymphoma viability. Thus, we treated cell lines of different lymphoma subtypes with the PL1 mAb, which recognizes the PSGL-1 N-terminal extracellular region and blocks functional interactions with selectins.^24^ Importantly, the PL1 mAb also detected surface PSGL-1 in T and B lymphoma cell lines (Fig. S1A). Stimulation of Sézary syndrome HUT-78 and mycosis fungoides HH cell lines with PL1 mAb for 24 h led to a strong decrease in cell viability (Fig. 2A). Next, we tested the effect of anti-PSGL-1 treatment in other lymphoma cell lines and found that the SU-DHL-1 (ALCL), OCI-LY3 (DLBCL), and U2932 (DLBCL) cell lines also had drastic reductions in cell viability (Fig. 2A). By treating the HUT-78 and U2932 cell lines with different PL1 dilutions, we observed that the killing action of PL1 was dose-dependent (Fig. S1B). To control for possible indirect toxic effects of the PL1 hybridoma preparation, we used as a control an unrelated hybridoma mAb from the same vendor (targeting PAX5 transcription factor), and found no induction of HUT-78 or U2932 cell death (Fig. S1C). Showing that the PL1-mediated killing effect was specific to malignant cells, PL1 treatment did not impair viability of healthy donor T cells (Fig. 2B). Finally, and showing that the antibody-mediated cell killing was not restricted to the PL1 mAb, we found that the KPL-1 mAb could also kill HUT-78 cells in a dose-dependent manner (Fig. S1D). These findings indicate that surface PSGL-1 antibody targeting can kill lymphoma cell lines of multiple subtypes, independently of PSGL-1 surface levels.

**Figure 2.**
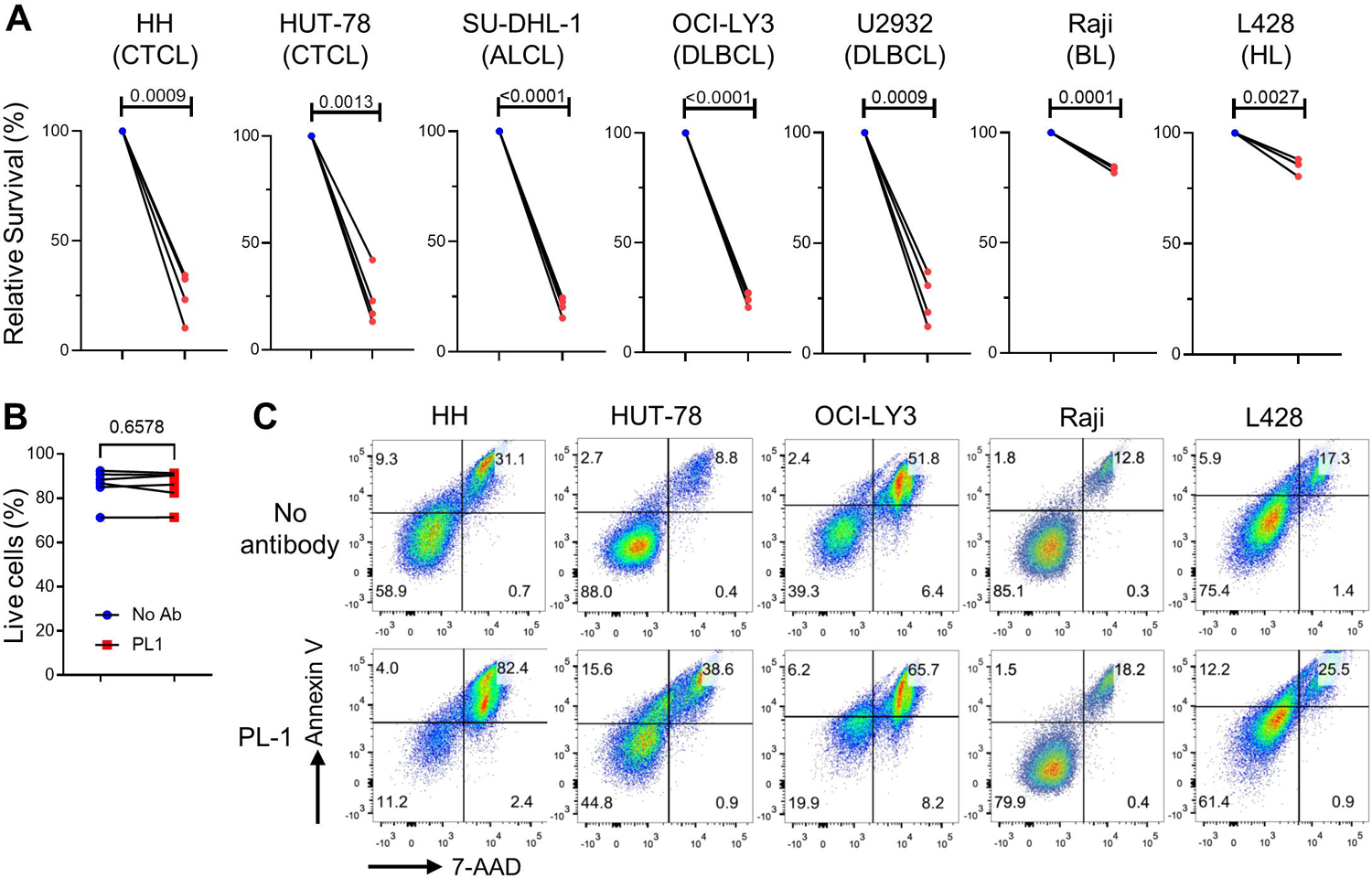
PL1 mAb treatment leads to lymphoma cell apoptosis. A) Viability of lymphoma cell lines after 24 h of culture with or without PL1 mAb. Plotted data represent four independent experiments performed. *P* values were determined by paired *t* tests. B) Percentage of viable healthy donor T cells after culture for 24 h with or without PL1 mAb. Plotted data represent values for six independent individuals. C) Flow cytometry plots of Annexin V and 7-AAD staining of lymphoma cell lines culture with or without PL1 mAb for 24 h.

To determine if the PSGL-1 mAb induced cell death though apoptosis, we analyzed the annexin V levels in cell lines treated for 24 h with PL1. Indeed, PL1 increased the frequency of annexin V and 7-AAD double-positive cells in HUT-78, HH and OCI-LY3 cell lines (Fig. 2C), thus indicating that the PL1-mediated reduction in cell viability was caused by apoptosis.

### Anti-PSGL-1 induces lymphoma cell apoptosis through the MAPK signaling pathway

To investigate the molecular mechanisms through which anti-PSGL-1 treatment was causing lymphoma cell death, we assessed whether the PL1 mAb could activate specific signaling pathways. Since the PL1 antibody was shown to activate ERK kinase in neutrophils,^43^ we studied ERK phosphorylation upon PL1 treatment of the PL1-sensitive HUT-78, OCI-LY3, U2932 cell lines and the PL1-resistant Raji cell line. ERK phosphorylation in HUT-78 cells was increased upon PL1 treatment, whilst no major differences were observed for other cell lines (Fig. 3A). To understand if the MAPK signaling pathway was involved in the PL1-induced cell death, we pre-treated lymphoma cell lines with a small molecule MEK1/2 inhibitor before stimulation with the PL1 mAb. As expected, we observed a significant reduction of phosphorylated ERK levels in HUT-78 cells upon MEK inhibition (Fig. S2). We next treated HUT-78, U2932 and OCI-LY3 cells with the MEK inhibitor for 8 h in the presence or absence of PL1 mAb. While the MEK1/2 inhibitor had no impact on HUT-78 and U2932 viability, it increased OCI-LY3 cell death (Fig. 3B). Strikingly, cell death induced by PL1 treatment was abrogated by the MEK1/2 inhibitor in HUT-78 and U2932 and to lower extent in OCI-LY3 cells (Fig. 3B). These data suggest that PL1 treatment induces apoptosis through activation of the MAPK signaling pathway.

**Figure 3.**
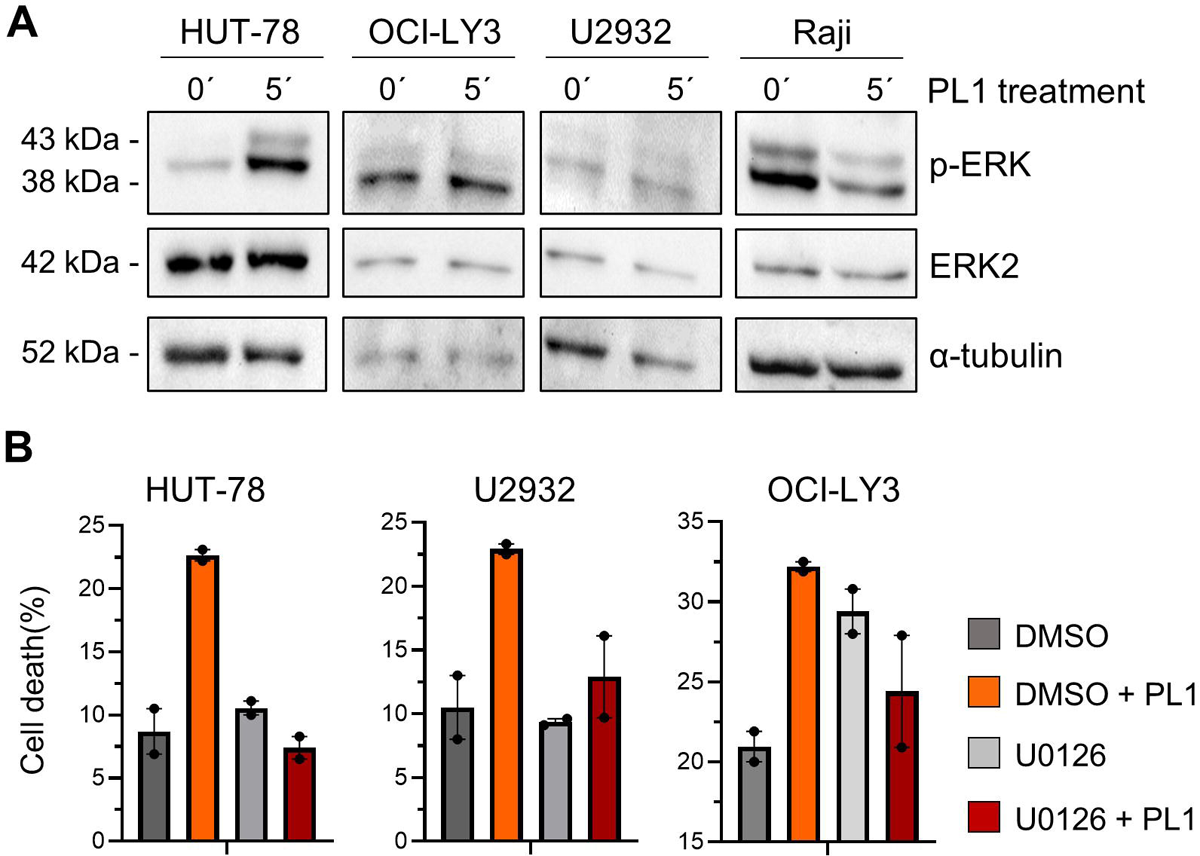
PL1-induced cell death is blocked by a MAPK inhibitor. A) Levels of phosphorylated ERK (p-ERK), ERK and α-tubulin in HUT-78, U2932, OCI-LY3 and Raji cell lines before and after 5 min of PL1 treatment. All cells were put under starvation conditions (only RPMI, no supplements) 2 h prior and during the experiment. B) Viability of HUT-78, U2932 and OCI-LY3 cell lines after 8 h of PL1 treatment in the presence or absence of 10 μM U0126 MEK1/2 inhibitor.

### *In vivo* administration of anti-PSGL-1 reduces cutaneous T-cell lymphoma growth

To determine whether the PL1 antibody could induce apoptosis *in vivo* and hamper tumor growth, we administered PL1 to immunodeficient mice xenografted with the PL1-sensitive HUT-78 cell line, which we beforehand confirmed to efficiently generate s.c. tumors (Fig. 4A).

**Figure 4.**
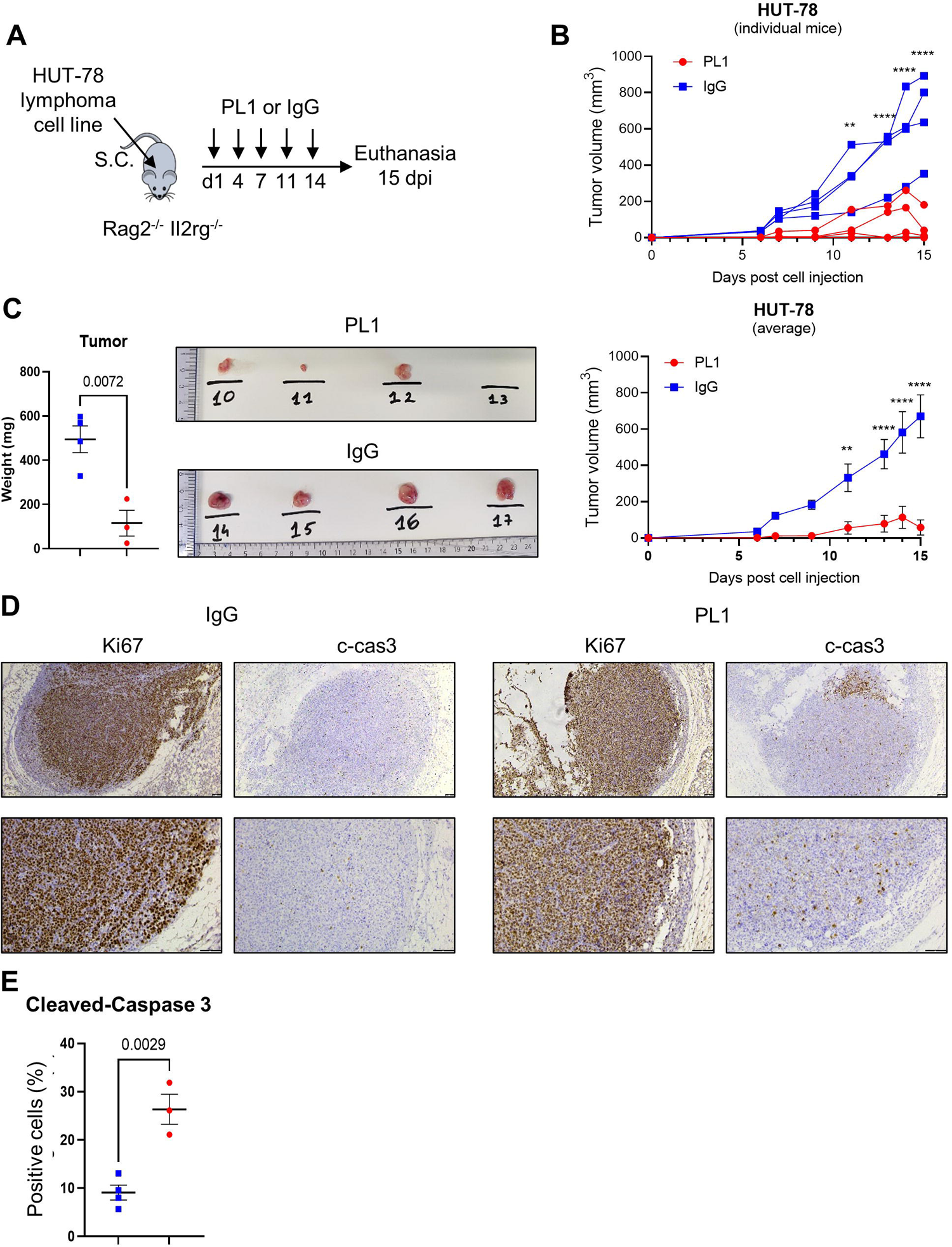
PL1 treatment significantly impairs HUT-78 tumor progression *in vivo* by promoting apoptosis. A) Schematic of the *in vivo* experimental design. B) Individual (top panel) and average (bottom panel) tumor volumes of HUT-78-injected mice treated with control IgG (n=4) or PL1 mAb (n=4). C) Tumor weights of HUT-78-injected mice at experimental endpoint. D) Representative immunohistochemistry of Ki67 and cleaved caspase 3 (c-cas3) staining of HUT-78-derived tumors from IgG- or PL1-treated mice. Scale bar,100 µm. E) Percentage of c-cas3-positive immunolabeled apoptotic cells in HUT-78-derived tumors from IgG-(n=4) or PL1-treated (n=3) mice. In panels B, D and E, *P* values were determined by unpaired *t* test.

Strikingly, HUT-78 tumor growth was significantly impaired upon PL1 treatment, as compared to IgG-treated mice (Fig. 4B). At the experimental endpoint, the tumor weight in PL1-treated mice was significantly decreased compared with IgG-treated mice (Fig. 4C). In the PL1 treatment group, tumors appeared in all 4 animals, but these regressed immediately with the continuous treatment (Fig-4B), with even complete tumor remission for one mouse (mouse #13; Fig. S3A). In this 15-day experiment no visceral organs in tumor-bearing mice were enlarged or displayed signs of tumor dissemination, with or without PL1 treatment (Fig. S3B). To verify if PL1-induced tumor inhibition was caused by a block in cell proliferation or increased apoptosis, tumors were immunostained for Ki67 and effector caspase 3 cleavage (c-cas3). Ki67 widespread immunostaining in PL1-treated and control tumors showed that PL1 did not arrest cell cycling (Fig. 4D). In contrast, the percentages of c-cas3-positive tumor cells were significantly increased in PL-1-treated mice compared with IgG controls (Fig. 4D and E). These results demonstrate that anti-PSGL-1 treatment induces CTCL apoptosis *in vivo* and substantiate the idea that PSGL-1 mAbs are of therapeutic value against this disease.

### Anti-PSGL-1 does not hamper Raji lymphoma growth

To rule out the possibility that anti-PSGL-1 targeting could impact tumor growth due to an indirect effect on innate immune cells present in *Rag2*^-/-^*Il2rg*^-/-^ recipient mice, we treated mice xenografted with Raji lymphoma cells with PL1 mAb for 14 days (Fig. 5A). The Raji s.c. tumor model progressed more slowly than the HUT-78 model (Fig. 5B), and was accompanied by mild splenic enlargement (Fig. 5D). In contrast to what was observed with the HUT-78 model, the PL1 mAb had no significant impact on Raji tumor progression (Fig. 5B). Furthermore, no significant differences in tumor and organ weight between PL1- and IgG-treated mice were observed at experimental endpoint (Fig. 5C and D). These results indicate that the PL1 mAb anti-tumoral function *in vivo* correlates with its effects *in vitro* and it was dependent on the direct action on tumor cells rather than indirect effects on innate immune responses.

**Figure 5.**
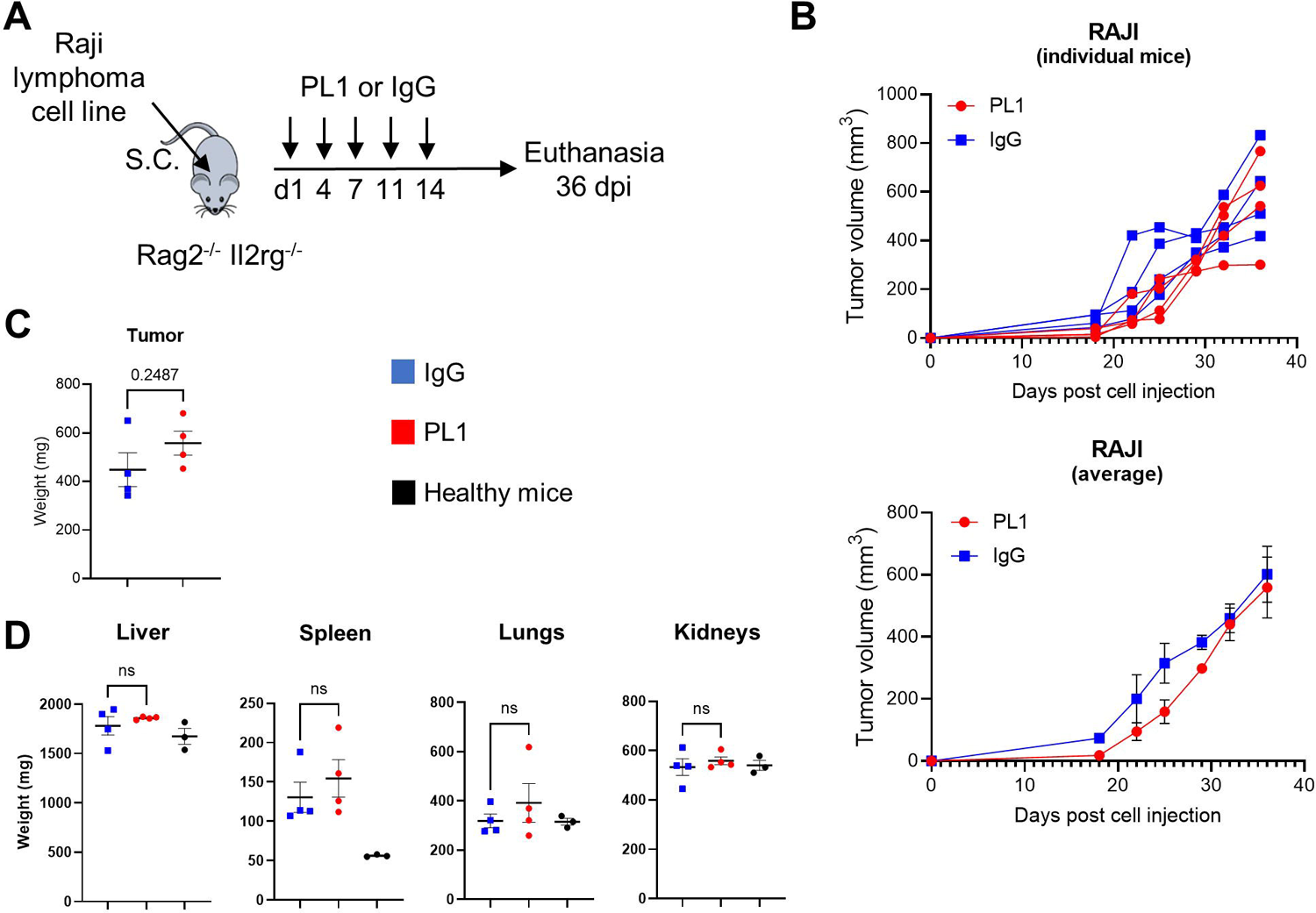
PL1 treatment does not impact Raji tumor development *in vivo*. A) Schematic of *in vivo* experimental design. B) Individual (top panel) and average (bottom panel) tumor volumes of Raji-injected mice treated with control IgG (n=4) or PL1 (n=4). C) Tumor weights of Raji-injected mice at experimental endpoint. D) Organ weights of Raji-injected mice (24 weeks) at experimental endpoint and non-injected *Rag2*^-/-^*Il2rg*^-/-^ mice. *P* values determined by unpaired *t* tests.

## Discussion

Here, we showed that PSGL-1 targeting may be a therapeutical approach in lymphoma. The PSGL-1 glycoprotein was expressed in most lymphoma cell lines analyzed and its antibody targeting resulted in *in vitro* killing of several cell lines by apoptosis. As a preclinical therapeutic proof-of-concept, we found that anti-PSGL-1 potently inhibited the HUT-78 CTCL cell line *in vivo* growth.

Two distinct antibodies targeting the N-terminal extracellular region of PSGL-1, PL1 and KPL-1, induced HUT-78 cell death. The two antibodies have overlapping human PSGL-1 epitopes, including amino acid residues 49-62 (LDYDFLPETEPPEM) for PL1 and residues 46-52 (YEYLDYD) for KPL-1, and both block PSGL-1 interactions with selectins.^44,45^ Therefore, we posit that both antibodies cause cell death through similar mechanisms. Previous research from Peru et al^46^ showed that pre-treatment of the CTCL My-La cell line with a mAb targeting CLA significantly impaired tumor growth in intrahepatic xenografted mice. Since CLA is a post-translationally modified variant of the PSGL-1 glycoprotein,^29^ our findings that two distinct anti-PSGL-1 mAbs killed HUT-78 cells *in vitro* and the PL1 mAb significantly delayed HUT-78 tumor growth in mice corroborates the notion that targeting the PSGL-1 glycoprotein extracellular region with antibodies has a therapeutic potential against lymphoma.

Although T-cell lymphoma cell lines were sensitive to PL1 treatment, normal healthy donor T cells were not killed by this mAb ligation, which suggests that specific molecular characteristics in these malignant cells render them vulnerable to PSGL-1-induced cell death. The intracellular activation status of lymphoid cells might play a role in PSGL-1 ligation induced cell death. It was reported that PSGL-1 antibody cross-linking induced apoptosis of mouse activated T cells.^47^ Since CTCL and ALCL cells usually present an activated phenotype,^48–51^ it is possible that apoptosis in these cells might result from combined signaling induced by PSGL-1 antibody ligation and constitutively active TCR signaling. Likewise, the B-cell lymphoma cell lines that were sensitive to PL1 are derived from activated B-cell-like DLBCL, so it is tempting to speculate that the activated status of these lymphoma cells may have rendered them vulnerable to PL1 treatment.

The molecular mechanisms through which the anti-PSGL-1 mAb causes lymphoma cell death remains unclear. It was proposed that the CLA mAb caused CTCL apoptosis by blocking its functions.^47^ The anti-PSGL-1 mAbs could act similarly or they could trigger directly pro-apoptotic signaling pathways. It was earlier reported that the PL1 mAb could not only block PSGL-1 interactions with selectins,^24^ but also trigger the activation of signaling proteins such as the Syk and ERK1/2 kinases.^43,52^ We found that PL1 ligation induces MAPK signaling pathway in the HUT-78 cell line through phosphorylation of ERK kinases. Moreover, inhibition of this signaling pathway hindered the apoptotic effect of PL1 in the HUT-78 and U2932 cell lines. Therefore, PL1 likely kills lymphoma cells through PSGL-1 ligation rather than PSGL-1 blockade.

Although our findings indicate that PSGL-1 antibody targeting can be a therapy for a broad range of lymphomas, it was also evident that not all cases are targetable by PL1 treatment. For example, the PSGL-1-expressing Raji and L-428 cell lines were poorly sensitive to PL1 treatment *in vitro* and this mAb had no significant effect on Raji tumor growth *in vivo*. Our *in vitro* results showed that the PSGL-1 surface expression levels per se did not strictly correlate with the PL1-mediated decrease in cell viability. For example, the U2932 and OCI-LY3 cell lines displayed lower levels of surface PSGL-1 than the Raji and L-428 cell lines, and yet the viability of the former was much more severely reduced by PL1 mAb than the latter. Although we cannot exclude that differences in PSGL-1 glycosylation status could explain the differential response to PL1 ligation, our data suggests that specific intracellular molecular mechanisms rather than PSGL-1 surface levels determine susceptibility to anti-PSGL-1-induced cell death. It was reported that the PL1 mAb binding to the PSGL-1 extracellular region depends on O-linked glycosylation,^24^ while the KPL1 mAb binding does not.^44^ The direct comparison of PL1 and KPL1 staining of five lymphoma cell lines showed that PL1 staining was generally lower than KPL1 staining (Figure S1A). The lower levels of PSGL-1 staining by PL1 could be due to the presence of non-glycosylated PSGL-1 protein at the surface of lymphoma cell lines. Although differential glycosylation could play a role in cell line susceptibility to PL1-induced cell death, the OCI-LY3 cell line had low PL1 staining, but yet was readily killed by this mAb. This suggests that PL1 can trigger cell death even at low levels of non-glycosylated PSGL-1.

Antibodies against the PSGL-1 or CLA glycoproteins appear to be both effective against CTCL. However, PSGL-1 is expressed by several myeloid and lymphoid malignancies such as acute myeloid leukemia, multiple myeloma, primary effusion lymphoma and ALCL,^16,17,20,36^ while CLA expression was observed in a single lymphoblastoid mantle cell lymphoma presenting skin lesions.^53^ The more widespread PSGL-1 expression than CLA expression and our finding that the PL1 mAb could kill CTCL, ALCL and DLBCL cell lines suggest that anti-PSGL-1 therapy might be broadened to a wider range of malignancies than anti-CLA therapy. In this line, Belmonte et al recently reported that PSGL-1 antibody treatment can induce ALCL cell death.^20^

Earlier research revealed that PSGL-1 promoted tumor growth and dissemination in mouse models of lymphoma and multiple myeloma.^36,39,40^ The finding that PSGL-1 antibodies trigger malignant lymphoid cell death indicates that the selective advantage of PSGL-1 expression for lymphoma or leukemia development, may render malignant cells vulnerable for targeted immunotherapy.

## Conclusion

We found that PSGL-1 is expressed in several lymphoma cell lines and its antibody targeting decreased their cell viability. Our research substantiates PSGL-1 as a promising therapeutic target in lymphoma. Further pre-clinical studies, especially in patient-derived xenografts, are warranted to validate the clinical potential of PSGL-1 targeted antibody therapies. The cytotoxic effect against PSGL-1 was not directly correlated with the cell surface expression of PSGL-1. Therefore, additional mechanistic studies are required to understand the PSGL-1 function different lymphoma types. This knowledge will pave the way for customizing this antibody therapy to each individual patient.

## Supporting information

Supplemental Figures and Raw Data

## Acknowledgments

We thank Alexandre M. Carmo, Ana X. Carvalho and Neil D. Perkins for providing cell lines. The authors acknowledge the support of the Animal Facility, Cell Culture and Genotyping, and Translational Cytometry i3S Scientific Platforms and the i3S Scientific Platform Histology and Electron Microscopy, member of the national infrastructure PPBI - Portuguese Platform of Bioimaging (PPBI-POCI-01-0145-FEDER-022122). This work was supported by Fundação para a Ciência e a Tecnologia (Portugal), European Social Fund, European Regional Development Fund (NORTE-01-0145-FEDER-000029, POCI-01-0145-FEDER-007274 and CEECINST/00091/2018/CP1500/CT0020 grants, IF/00056/2012 contract to NRdS and SFRH/BD/147979/2019 fellowship to J.L.P.), and grants from Gilead Sciences Portugal (Programa Gilead GÉNESE PGG/038/2017), and Associação Portuguesa Contra a Leucemia (APCL) in partnership with Sociedade Portuguesa de Hematologia (SPH) and Gilead Sciences.

## Data availability statement

The data that support the findings of this study are available from the corresponding author upon reasonable request.

