## Supplemental Figures and Raw Data for "Antibody targeting of surface PSGL-1 glycoprotein leads to lymphoma apoptosis and tumorigenesis inhibition"

### Supplementary Figures

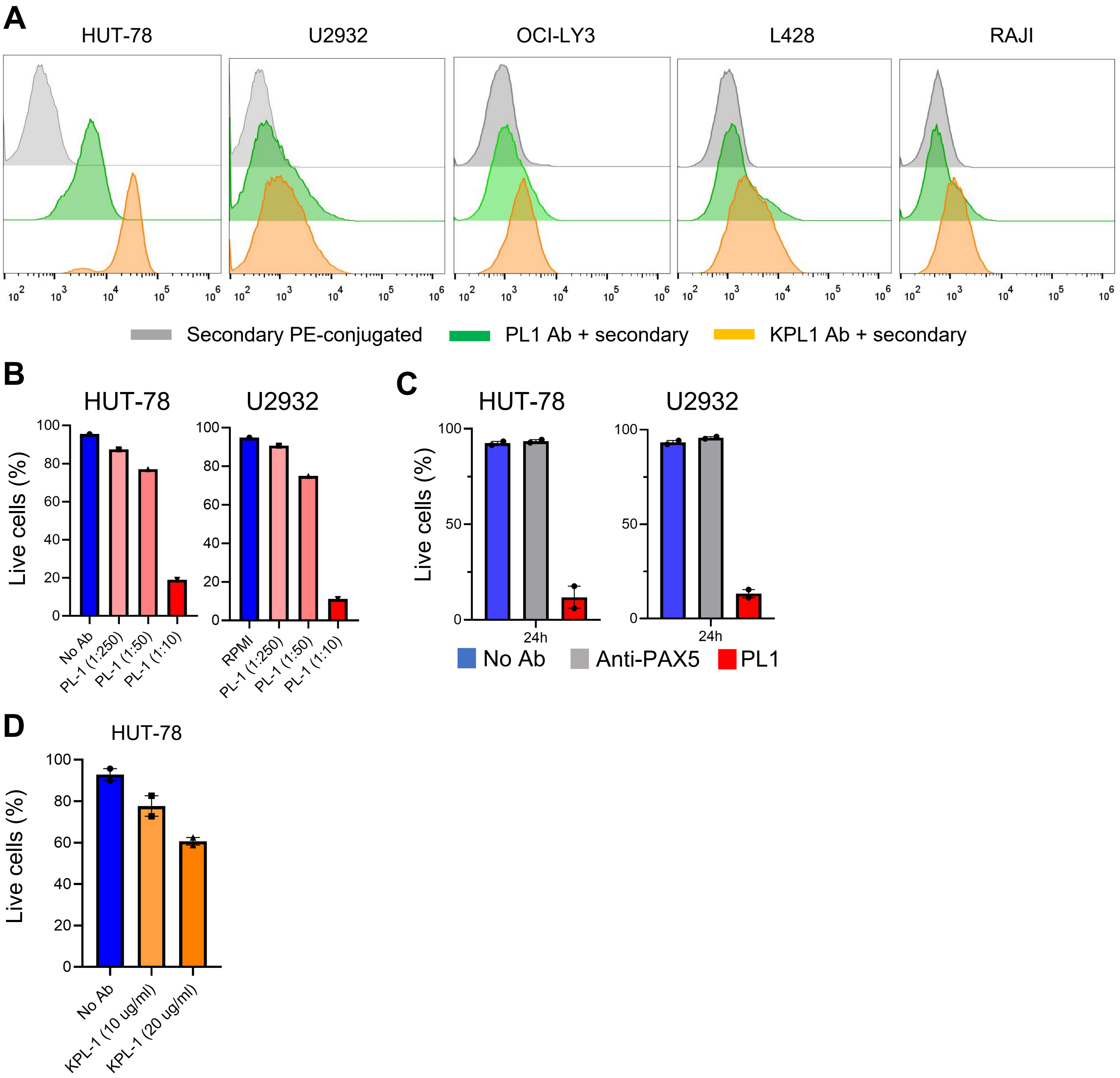


**Supplementary Figure 1. PL1 and KPL-1 clones detect PSGL-1 expression and induce lymphoma cell death.** A) Flow cytometry of PSGL-1 surface expression of HUT-78, U2932, OCI-LY3, L428 and Raji cell lines using PL1 or KPL-1 primary antibodies followed by PE-conjugated secondary antibody. B) Viability of HUT-78 and U2932 cell lines upon 24 h of culture with or without PL1 mAb, at the indicated PL1 dilutions. C) Viability of HUT-78 and U2932 cell lines upon 24 hours of culture with or without PL1 mAb and anti-PAX5 (PCRP-PAX5-1B7) mAb as control. D) Viability of HUT-78 cell line upon 24 h of culture with or without KPL-1 mAb, at the indicated concentrations.

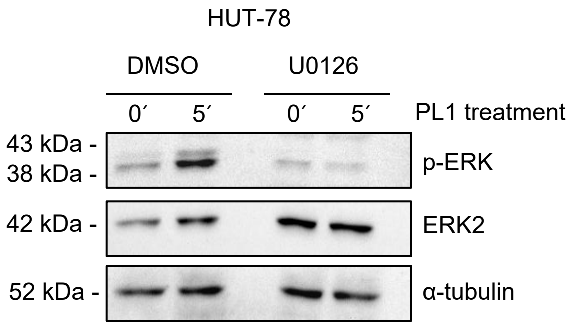


**Supplementary Figure 2. U0126 MAPK inhibitor blocks PL1-induced ERK phosphorylation in the HUT-78 cell line.** A) Levels of phosphorylated ERK (p-ERK), ERK and α-tubulin in the HUT-78 cell line with or without 10 μM U0126 treatment upon 5 min stimulation with PL1.

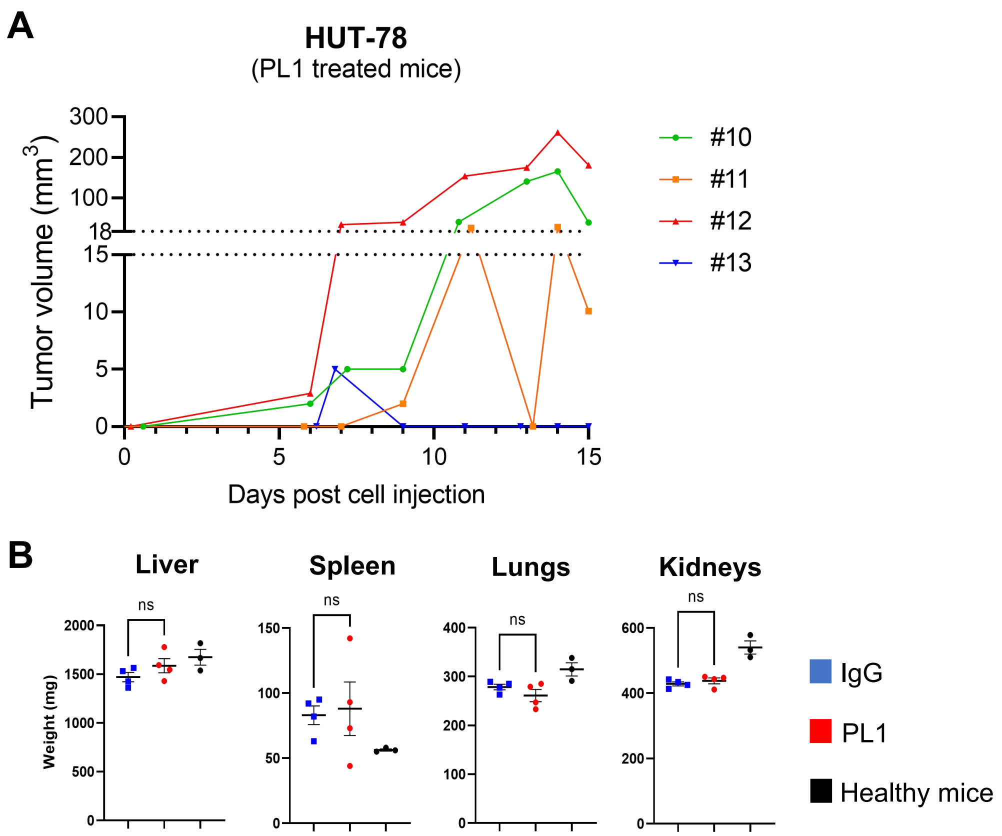


**Supplementary Figure 3. Tumor growth of PL1-treated mice.** A) Representation of HUT-78 tumor growth in four individual mice treated with PL1. B) Organ weight of HUT-78-injected mice at experimental endpoint and non-injected Rag2$^{-/-}$Il2rg$^{-/-}$ mice of similar age (13/14 weeks).

### RAW data

Original western blots, related to Figure 3A

P-ERK ERK2 Alpha-tubulin

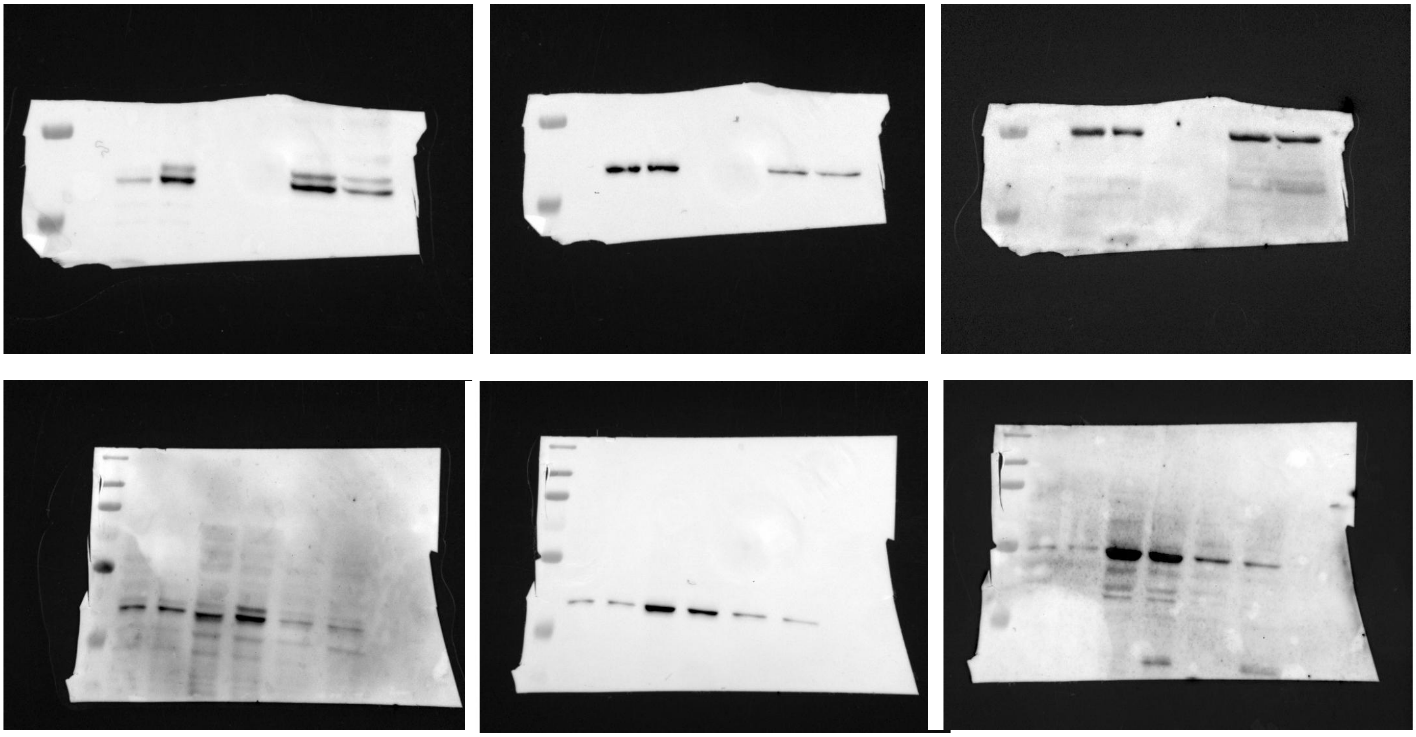

Original western blots, related to Figure S2A

P-ERK ERK2 Alpha-tubulin

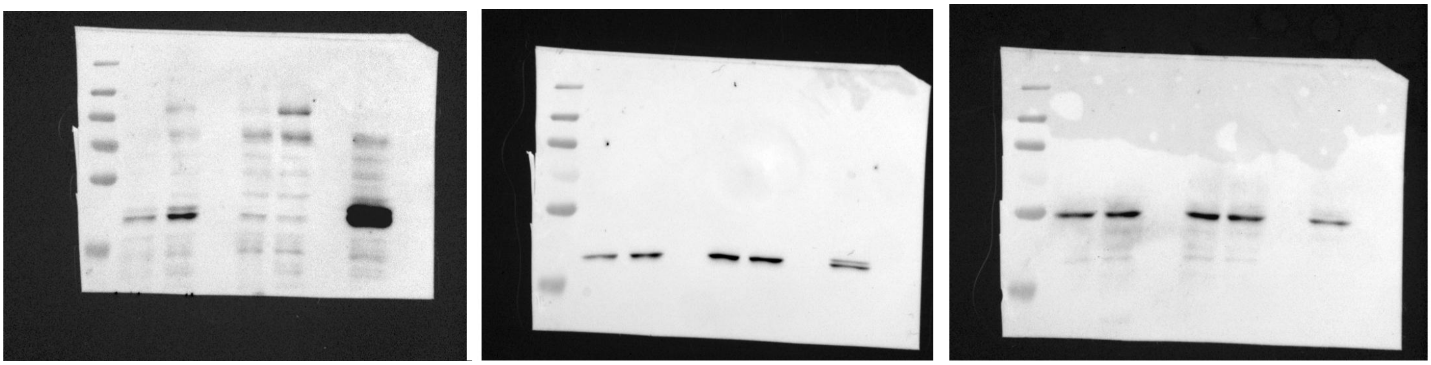
